# Microwell array based opto-electrochemical detections revealing co-adaptation of rheological properties and oxygen metabolism in budding yeast

**DOI:** 10.1101/2021.01.07.425712

**Authors:** Venkata Suresh Vajrala, Baptiste Alric, Adrian Laborde, Camille Colin, Emmanuel Suraniti, Pierre Temple-Boyer, Stephane Arbault, Morgan Delarue, Jérome Launay

**Affiliations:** CNRS, LAAS, 7 avenue du colonel Roche, F-31400, Toulouse, France; Université de Toulouse, UPS, LAAS, F-31400 Toulouse, France; Univ. Bordeaux, ISM, CNRS UMR 5255, INP Bordeaux, Pessac, France

**Keywords:** microdevices, microwells, ring nanoelectrodes, cyclic voltammetry, GEMs single-particle tracking, yeast cells, oxygen consumption, bioenergetics

## Abstract

Microdevices composed of microwell arrays integrating nanoelectrodes (OptoElecWell) were developed to achieve dual high-resolution optical and electrochemical detections on single *Saccharomyces cerevisiae* budding yeast cells. Each array consists in 1.6 × 10^5^ microwells of 8 µm diameter and 5 µm height, with a platinum nanoring electrode for in-situ electrochemistry, all integrated on a transparent thin wafer for further high-resolution live-cell imaging. After optimizing the filling rate, 32% of cells were effectively trapped within microwells. This allowed to analyse *S. cerevisiae* metabolisms associated with basal respiration while simultaneously measuring optically other cellular parameters. In this study, we focused on the impact of glucose concentration on respiration and intracellular rheology. We found that while oxygen uptake rate decreased with increasing glucose concentration, diffusion of tracer nanoparticles increased. Our OptoElecWell based respiration methodology provided similar results compared to the commercial gold-standard Seahorse XF analyser, while using 20 times lesser biological samples, paving the way to achieve single cell metabolomics. In addition, it facilitates an optical route to monitor the contents within single cells. The proposed device, in combination with the dual detection analysis, opens up new avenues for measuring cellular metabolism, and relating it to various cellular physiological and rheological indicators at single cell level.

## Introduction

Many physiological and molecular functions happening in cells are the sum of sequential processes, where nearly every biochemical reaction is a reaction–diffusion process involving the diffusive flux of biomolecules to cellular active sites (Kinsey et al., 2011). Unlike usual soft materials, the cells deal with non-equilibrium active forces and are able to transform cellular external cues into biochemical signals (Mizuno et al., 2007; Tee et al., 2009; Trepat et al., 2007). On the contrary, cells are capable of using biochemical signals to modify their viscoelastic properties, not only at the plasma membrane level, but also around the nucleus, within the intracellular organelles and the cytoplasm (Abidine et al., 2018; Kim et al., 2018; Makhija et al., 2016; Mathieu and Manneville, 2019). As such, the interplay between the cellular rheological and metabolic properties appear extremely important in overall functioning, through numerous pathways, in response to changes of their physico-chemical environment. For example, the regulation of ATP, cellular energy derived from digested glucose and consumed oxygen via mitochondrial oxidative phosphorylation OXPHOS, is linked to cellular membrane stiffness fluctuations and other cellular rheological changes (Alibert et al., 2017; Betz et al., 2009; Guo et al., 2014; Weber et al., 2012). In this regard, there has been a great deal of interest to understand how the rates of cellular respiratory metabolism (OXPHOS activity) is related to the intracellular rheological parameters and molecular diffusion within the cells. In order to have a comprehensive knowledge on such dynamic relationships, there is a need for integrated multi-parametric detection systems that can offer simultaneous monitoring of biochemical and biophysical properties of cells. The main focus of the presented research work concerns a novel dual-detection approach for simultaneous monitoring of average cellular oxygen consumption rate (OCR) and cellular rheological properties at single cell level.

Several electrochemical and optical sensor platforms are currently available for cellular respiration monitoring OCR, such as Clark-type electrodes in suspended biological samples (Kahn, 1964; Oroboros Instruments), or fluorescent probes in plated cultured cells (MitoXpress Xtra; Seahorse XF). In presence of specific drugs (metabolic uncouplers and inhibitors), oxygen consumption experiments can determine basal and maximal cellular respiratory capacity and the estimation of substrate effects, among other parameters (Simonnet et al., 2014; Wu et al., 2007; Zhang et al., 2012). Electrochemical methods, in particular ultramicroelectrodes (UME) and/or nanoelectrodes, can sense the concentration gradients of redox species up to few pico-litre volume range (Amatore et al., 2008; Cox and Zhang, 2012; Díaz-Cruz et al., 2020; Ino et al., 2017). The diffusion volume probed by a nanoelectrode would indeed be adequate for sensing the microenvironment of a single cell, where they offer steady-state diffusion limited currents and short response times (Amatore et al., 2010, 2006; Cox and Zhang, 2012; Date et al., 2011; Nebel et al., 2013; Santos et al., 2017). However, this approach is still been faced with challenges in dealing with the low throughput owing to the manual positioning of the electrode near the cell and UME spatial coverage over the cell, i.e., partial detection of targeted cellular events due to lateral diffusion phenomenon. To overcome these limitations, we have recently developed a microwell array integrated with recessed ring-type micro/nanoelectrodes (RME/RNE) (Sékli Belaïdi et al., 2016; Vajrala et al., 2019). A functional and intricate integration of these RNE-based devices within microwells allowed us to entrap individual mitochondria in a high-throughput manner, and followed their respiration under several metabolic activation and inhibition conditions.

In parallel, high-resolution fluorescence based single particle tracking (SPT) techniques have been widely used in the field of cell biology as a means of unravelling diffusion dynamics of biomolecules and tracers within cells, at nanometric spatial resolution (Kusumi et al., 2005; Mandal et al., 2016; Wirtz, 2009). Many fluorescent probes are available right now, where typically the probe is attached to the molecule of interest, such as an organelle, virus-like particles or a receptor to track the individual dynamics (Alcor et al., 2009; Rose et al., 2020). These tracer nanoparticles move within the cells and enable the study of cellular biophysical properties under different dynamic environments. Various groups have studied the motion of non-biological nanoparticles in cells, but typically, these techniques are labour-intensive and perturb the cellular natural behaviour due to their non-biocompatibility or bulkier sizes. We recently developed genetically encoded multimeric nanoparticles (GEMs) which are bright 40nm tracer particles of a defined shape, and relatively impervious to specific interactions with in the cell (Delarue et al., 2018, 2017). Using this novel methodology, we are able to measure the rheological parameters of a cell at the single cell level and without the need to inject fluorescent probes.

In this work, we developed a novel microwell array device which allows to intimately blend the electrochemical and high-resolution optical sensing techniques, to achieve dual-detection of entrapped cells, in an orderly fashion, and their responses over time. Such systems can allow simultaneous dual optical and electrochemical measurements to shed light on the intricate cross-relationships between cellular internal dynamics of biomolecular diffusion and OXPHOS based energy metabolism. We present a new generation of very well ordered microwell array device (OptoElecWell) with dimensions optimized for the study of the model fungus *Saccharomyces cerevisiae*, wherein the microwells (8 µm ***d*** × 5 µm ***h***) are connected to Pt nano-ring type working electrode, supported by a very thin (≈ 170 nm) glass substrate, used as a platform for monitoring both cellular oxygen consumption and high-resolution nanoparticle tracking at single cell level.

We first fabricated the microwell array chip, made up of “glass (170µm) /SiO_2_ (2.5µm) /Ti (20nm) /Pt (200nm) /Ti (20nm) /SiO_2_ (2.5µm)” sandwich stack using chemical vapour deposition, reactive ion etching and glass polishing techniques. Subsequently, the optical and electrochemical characterization of the microwell array were performed to realize the best performing conditions for, (i) Pt nano-ring working electrode, (ii) filling of wells with yeast cells, and (iii) nanoparticle tracking within yeast cells entrapped within microwells. Later, we demonstrated the integrated dual-detection approach for simultaneous electroanalysis of cellular oxygen consumption and fluorescence-based single particle diffusion measurements within cells. The simultaneous SPT and OCR measurements led to the counter-intuitive discovery of a co-adaptation of OXPHOS and rheological properties: as glucose concentration decreases, OXPHOS and supposedly ATP production increases while diffusion decreases.

## Materials and methods

### Preparation of yeast cells and GEMs expression

Budding yeast *(Saccharomyces cerevisiae*, derived from BY4742 strain) cells, labelled on their histone by a mCherry tag to visualize the nucleus and express genetically encoded multimeric nanoparticles - GEMs (Delarue et al., 2018) (courtesy of L. Holt) were cultured on synthetic complete medium agar. They were resuspended 24h before experiments in liquid medium with different glucose concentration (0.5 g/mL, 1 g/mL and 2 g/mL), and when necessary were diluted in fresh buffer to 0.5 OD (optical density) at the beginning of the measurement.

### Oxygen consumption rates measurements using Seahorse XF bioanalyzer

Yeast (*S. cerevisiae*) oxygen consumption rates were measured using a SeaHorse XF 24 Analyzer (Seahorse Biosciences, USA). The experiment was performed in yeast cell culture solution containing 1.5 × 10^5^ cells per well of 24-well plates (600 µL). Basal respiration conditions were measured for cells in the presence of three extracellular glucose concentrations, 0.5%, 1% and 2% (w/v).

### Microfabrication of OptoElecWell device

The design and fabrication of the microwell array reported here is inspired from our previous works where we reported the first generation of OptoElecWell and discussed its design by multiphysics simulation, manufacturing process and characterization (Sékli Belaïdi et al., 2016; Vajrala et al., 2019b).

Photo-mask designs for the microwell arrays and electrodes (e1 and e2), as shown in figure 1b-c, were first drawn in CleWin software and later printed on high-resolution glass substrates. Starting from B33 glass substrate (500 µm thick), three deposition steps were conducted in a row, the first step being 2.5 µm thick SiO_2_ film by plasma enhanced chemical vapour deposition (PECVD). This step was followed by the deposition of 200 nm platinum layer by evaporation while using two 20 nm titanium interfacial layers, and another 2.5 µm thick PECVD SiO_2_ layer to form a SiO_2_/Ti/Pt/Ti/SiO_2_ stack. Next, the microwell array with integrated Pt-nanorings was created, through a patterned photoresist mask, where the whole stack (SiO_2_/Ti/Pt/Ti/SiO_2_) was etched using inductively coupled plasma reactive ion etching (ICP-RIE) with three different gaseous mixtures (CF_4_/Ar, SF_6_ and Cl_2_). Later, the electrical contacts of working electrodes e1 and e2 were established by exposing the platinum layer at selected areas using wet etching of SiO_2_. Lastly, in order to facilitate high-resolution optical measurements, the bottom surface of the B33 glass wafer was carefully polished using Logitech PM5 lapper, where the wafer was lapped in between two counter-rotating cast plates using slurries consisting of different alumina abrasive grains with defined size distribution. First, the wafer thickness was reduced from 500 µm to 300 µm using alumina slurry with 20 µm grain size for 60 minutes. Later, the wafer was carefully polished with a slurry of 9 µm grain size, followed by another lapping step with 1 µm grain sized slurry, to obtain 170 µm thick wafer. Finally, the electrical contacts of e1 and e2 for performing electrochemistry were established by micro-soldering technique using sliver epoxy silver paint.

**Figure 1:**
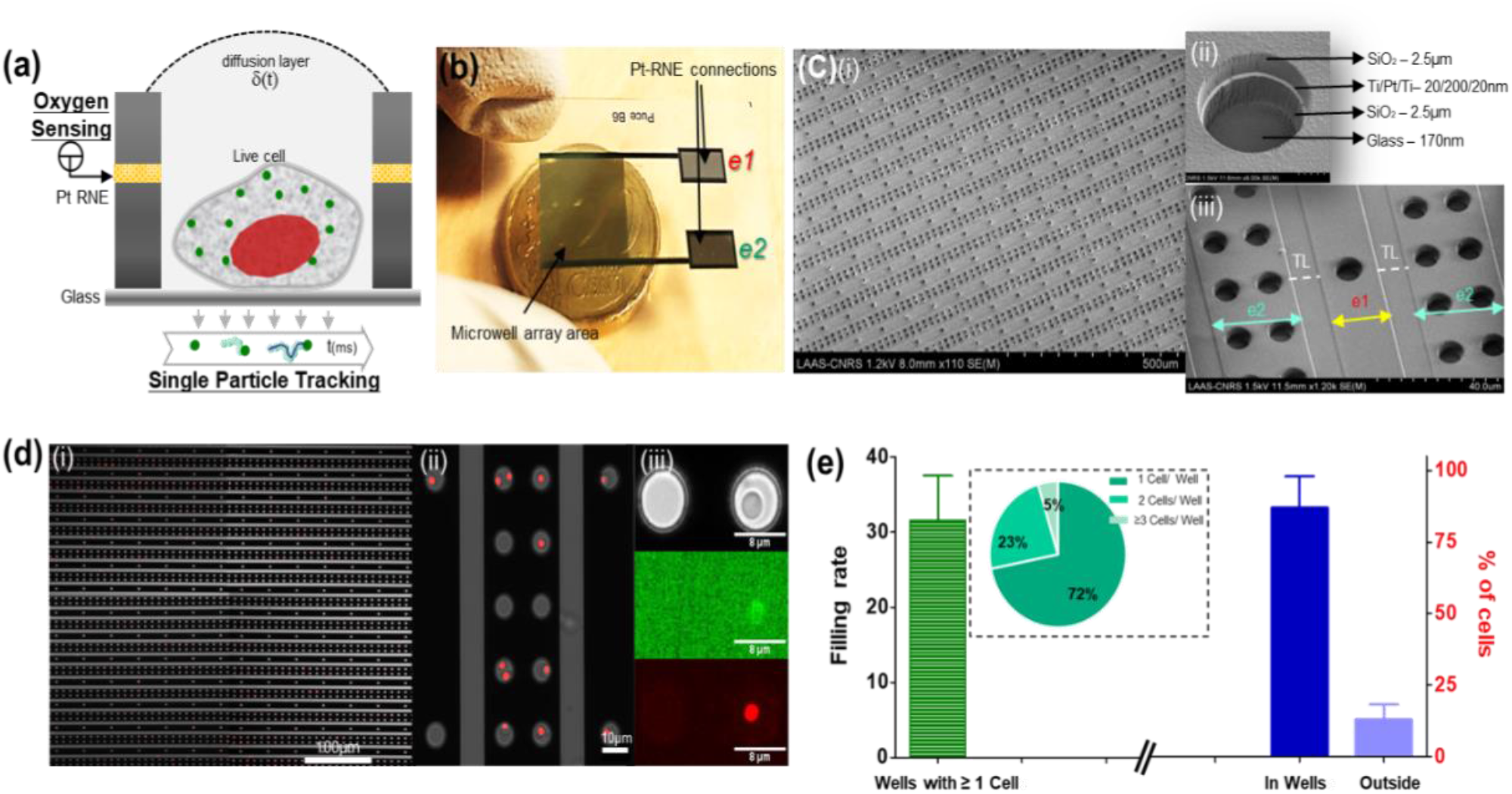
(a) Schematic view of the opto-electrochemical (dual) detection using OptoElecWell array to monitor yeast cellular metabolic responses; (b) Photograph of the OptoElecWell chip having 1.6 × 10^5^ microwells in comparison with a 20-cent coin. Pt-RNE means Platinum ring type nanoelectrodes; (c) Electron microscopic images of the device, showing microwell arrays (e1 and e2) with a mean well dimensions of 8 µm ***d*** and 5 µm ***h***, on a thin glass substrate having thickness around 170 nm. e1 and e2 represents the columns (subsets) of microwells within the array having 60 µm and 15 µm inter-well array distance respectively. In between the columns there exists a transparent layer (TL). (d) (i) The representative micrograph (overlay of a bright field and a fluorescence image in red) of yeast cells entrapped in OptoElecWells array after seeding with cells that were stained with nucleus tag (histone tagged with mCherry fluorophore); (ii) Zoom-in the area from (i), showing the trapped singlet and multiplet of cells in each microwell; (iii) Microscopic pictures of yeast cells showing nucleus and 40 nm-GEMs expressed in a single yeast cell within a microwell. The images were taken through the transparent microwell array chip using inverted fluorescence microscope at 5× and 20× magnifications; (e) The green bar (left) depicting the percentage of the filling rate within microwell array. The Pie chart represents the measured distribution of cell occupancy per well. The blue coloured bar graph (right) shows the percentage of cells found within the microwells vs outside the wells.

### On-chip dual detection of yeast cell metabolism

A plastic fluidic prototype system, supported by plastic screws and smooth silicone polymer pad was fabricated using micromilling process, to accommodate thin OptoElecWell array at the bottom. Yeast cells were immediately used for sample preparation after reaching the required OD. Just prior to use, the microwell array was cleaned carefully with DI water and ethanol, followed by oxygen plasma treatment (Diener plasma 50W) for 2 minutes. The microwell array was then fixed at the bottom of fluidic system and 150 µL of yeast cells were added to the reservoir. Yeast cells were gently moved back and forth using pipette on microwell array surface, over 30 cycles, followed by a 10 minutes incubation to promote the sedimentation of cells in microwells. Later, the leftover cell suspension was removed by performing careful washings. Finally, 1 mL of aerated yeast buffer was injected into the fluid reservoir, and all three-electrode connections, along with the microscopy focus was established to analyse the cells in microwells.

### Live cell imaging and single particle tracking

All the experiments were carried out at 30°C, on the stage of an inverted confocal microscope (Leica DMI 6000, Germany). Imaging was recorded on an scMOS camera (Hamamatsu). Cell nuclei were monitored by taking pictures using mCherry fluorescence (laser at 561 nm wavelength). We targeted the intrinsic T-sapphire fluorescent protein (green) to image 40nm-GEMs at 488 nm wavelength and sampled at a rate of one image every 20 milliseconds. The tracking of individual GEM nanoparticles was performed with the Mosaic suite of FIJI assuming Brownian dynamics and analyse on home-made programs on Matlab, available on demand (Delarue et al., 2018).

### Electrochemical characterization and oxygen consumption monitoring

Electrochemical measurements were conducted at 30°C, using Ag/AgCl reference (RE) and Pt-mesh counter electrodes (CE) connected to BioLogic potentiostat, equipped with ultralow current module. In this work, we used nanoelectrode array (e1) as the working electrode (WE) (figure 1b-c). The electrochemical activation and characterization of nano-ring type, sandwiched Pt layer was done by performing a series of cyclic voltammograms CVs in 1 M H_2_SO_4_ at 1 V/sec. We used continuous CV technique to monitor the cellular oxygen consumption at different metabolic stages (4 V/sec, 20 seconds of relaxation gap between each scan). Carbonyl cyanide-p-trifluoromethoxyphenylhydrazone (FCCP, Sigma-Aldrich) and antimycin A (AMA, Sigma-Aldrich) solutions were prepared in the sample buffer at the following final concentrations: FCCP 1 µM and AMA 10 µM. The reductive current values of each scan at −0.7 V were recorded and processed by a Python program. Origin software was used to plot the data and perform statistical analyses.

### Model for Oxygen consumption rate monitoring within OptoElecWell device

We assumed that the oxygen consumption of a cell and the oxygen reduction on a recessed disk microelectrode (Sékli Belaïdi et al., 2016) are similar. They both rely on the diffusion and concentration of oxygen available in the buffer, and according to the diffusion law applied to electrochemistry, the dissolved oxygen flux *ϕ* from the bulk solution to the cell surface is given by:

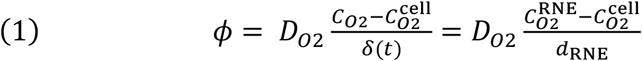

Where *D*_02_ is the diffusion coefficient of dissolved oxygen, 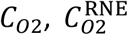 and 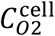 are the dissolved oxygen concentrations in the bulk solution, at the ring nanoelectrode surface and at the cell surface, *d*_RNE_ is the distance between the ring nanoelectrode and the cell, and *δ*(*t*) is the time-dependent diffusion layer thickness:

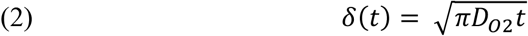

According to equation (1) and (2), 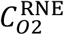 can be rewritten as:

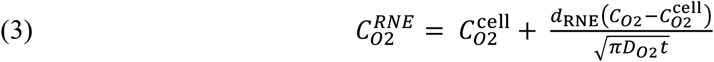

In equation (3), it should be noticed that 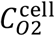 is representative of the cellular oxygen consumption and it depends on the glucose concentration in solution (Barnabé and Butler, 2000; Takeda et al., 2015). Since the dissolved oxygen reductive current *i*_RNE_ measured on the Pt ring nanoelectrode is proportional to 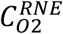 and the OptoElecWell device was designed to prevent crosstalk between microwells (Lemercier et al., 2017; Sékli Belaïdi et al., 2016), the measured currents should follow temporal variations according to a “Cottrell-like relation” (A.J. Bard and L R Faulkner, 2000).

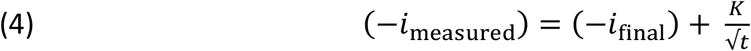

Where the final current *i*_final_ and the parameter *K*, directly proportional to the oxygen consumption rate, depends on the glucose concentration in solution.

## Results and discussion

Over the years, we have designed different ring-type nanoelectrode (RNE) based microwell arrays (Lemercier et al., 2016; Sékli Belaïdi et al., 2016; Vajrala et al., 2019b). We reported that the quality of electrochemical response of these arrays, which is the sum of electrochemical response of individual nanoring type microelectrodes, heavily relies on the distance between each microelectrode within the array and the number of microwells with biological species, i.e., active wells. In line with the dual-detection of cells, a nice compromise should be established so that the wells cannot be very close to each other (< 5 × R), which further increases the crosstalk between each microwell, or not so far from each other (> 20 × R) so that it decreases the number of wells having cells and further impacts the optical throughput. In addition, few cells can also be located at the inter-well surface area, which are optically untraceable, preventing the normalization and further complicating the sensing and metabolic interpretations. With this in mind and by considering all the constraints, we developed a novel microwell array (OptoElecWell) device (figure 1), where 1.6 × 10^5^ microwells having 8 µm diameter, 5 µm height, were arranged in a specific pattern such that there exists two different RNE well array zones, e1 and e2 with 60 µm and 15 µm interwell distances respectively (figure 1b-c). They were designed for the purpose of either connecting both RNEs together or individually addressing each electrode array to notably follow two electroactive species. In addition, we incorporated a transparent layer (TL, figure 1) in between the columns of microwells to estimate the number of cells that are not entrapped in the wells.

### Filling rate, monitoring and evaluation of yeast cells within OptoElecWell

The yeast cell array was prepared by dispersing the cell suspension in a fluid chamber that was carefully fitted on top of microwell array device. Oxygen plasma treated flat silicon surfaces favours fair immobilization of biological material, owing to the electrostatic interactions. Another advantage of plasma is that it remains a very simple surface modification strategy, where it enables the measurement of natural cell responses without the interference caused by complicated surface immobilization strategies. In this regard, we activated the OptoElecWell by treating the surface of array with oxygen plasma prior to cell injection. Cells were later loaded by dispersing a suspension of freshly cultured yeast cells and followed by careful washings. We used suspensions of 0.5 OD and observed that the cell concentration and washings could clearly affect the cell packing density (results not shown). Under optimized conditions, there were almost 32% of wells occupied by 1 or more cells and 75% within them were containing only one cell (figure 1d-e). From both optical and electrochemical monitoring point of view, it is very important to know the total number cells, along with the distribution of cells per well. Additionally, the electrochemical monitoring does not only rely on the filling rate but also the estimation of interfering cells that are sedimented at the inter-well surface (outside the wells); the lesser the better. We could estimate the percentage of total cells that were within the wells, being 87%, compared to the 13% of cells that were sedimented on the inter-wells surface area.

### Electrochemical characterization and monitoring of cellular oxygen consumption

We used in this study the electrode array (e1) having 1.7 × 10^4^ microwells in total as a working electrode. We chose e1, with the interwell distance of 60 µm such that there will be minimal overlapping of diffusion layers with neighbouring microelectrodes. The microwells with interwell distance of 15 µm (e2) were kept electrically neutral and used for optical monitoring of cells, to increase the optical throughput. The electrochemical performance of the nanoring-type, 200 nm thick platinum within the microwell array was first characterized by cyclic voltammetry in 1 M H_2_SO_4_ solution (potential scanned between −0.3 V to 1.6 V), showing clear proton reduction and oxidation waves similar to the native solid Pt metal (figure 2a). As a sampling method to sense local oxygen changes around the entrapped cells, within microwells, we chose continuous cyclic voltammetry over chronoamperometry. We noticed that under optimized conditions – higher scan rates and gaps between each scan (4 V/s and 20 s gap), CV allows to monitor the local chemical changes happening within the microwells, while allowing the oxygen concentration to get to the equilibrium before next measurement (Vajrala et al., 2019) i.e., no competition between working electrode and yeast cells in terms of oxygen usage. We first investigated the oxygen sensing capacity of OptoElecWells by performing the electrochemical reduction of dissolved oxygen under different oxygen concentrations in yeast buffer using cyclic voltammetry at a scan rate of 4 V/sec. The oxygen reduction peak current was detected around −0.7 V and demonstrated that the Pt nanoring electrode is sensitive to the local oxygen concentration changes (figure 2b).

**Figure 2:**
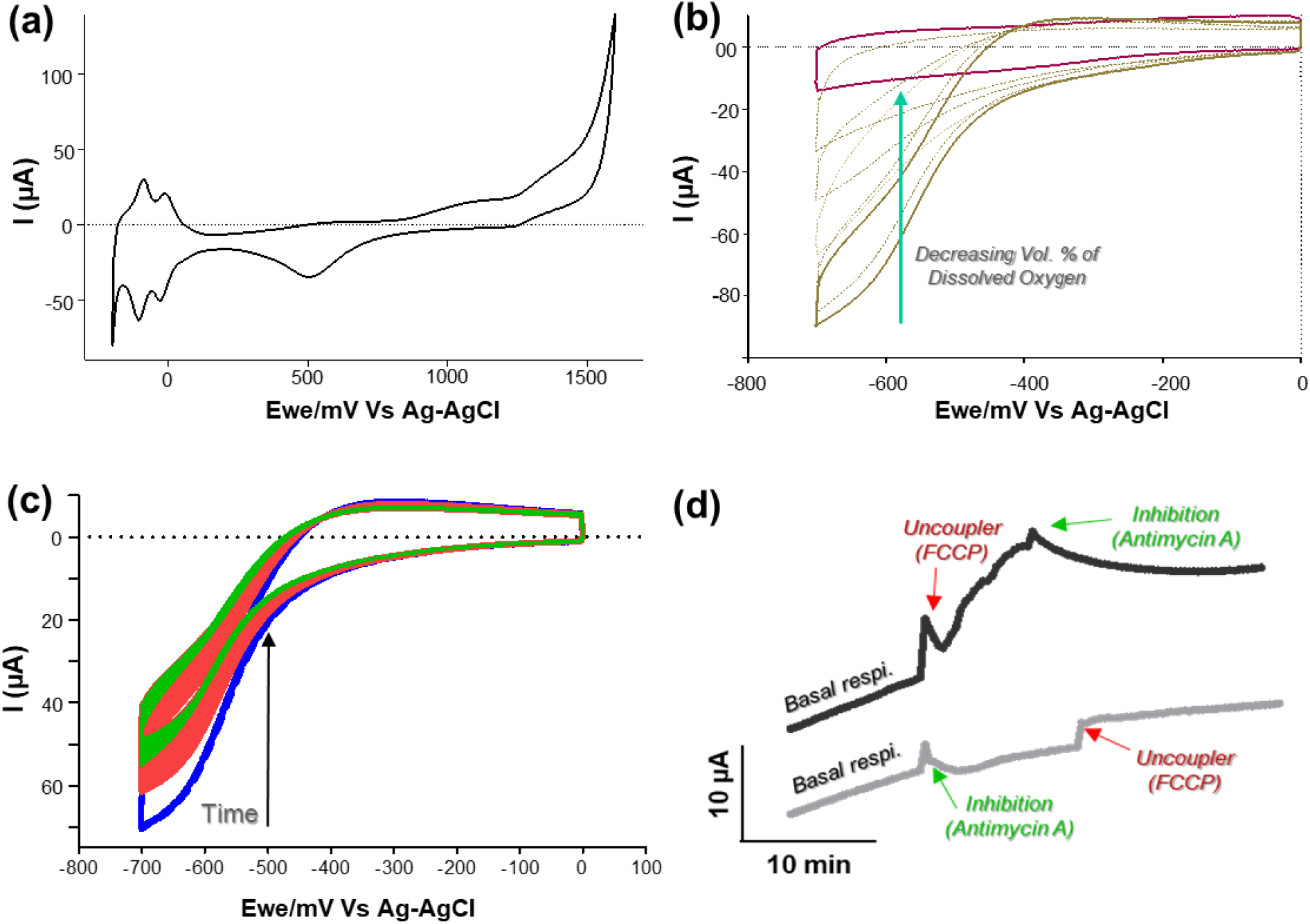
Electrochemical characterizations by cyclic voltammetry (CV) of the nanoelectrode based OptoElecWell device: (a) voltammogram (1 V/s) recorded in 1 M H_2_SO_4_ solution; (b) voltammograms (4 V/s) recorded in yeast buffer (pH 7.4) containing different oxygen concentrations; (C) A continuous series of cyclic voltammograms depicting the electrochemical monitoring of the oxygen consumption by yeast cells under different metabolic states. (Scan rate: 4 V/s, delay of 20 s between scans); (d) Oxygen reductive peak current evolution biased at −0.7 V vs Ag/AgCl measured on a continuous series of CVs (figure 2c), depicting the electrochemical monitoring of the oxygen consumption by yeast cells, followed by the additions of 1 µM FCCP and 10 µM Antimycin A. Ag/AgCl reference electrode and a platinum mesh as a counter electrode were used here.

Next, we entrapped budding yeast cells (*S. cerevisiae*) with a typical diameter of 8 μm within microwells, as a biological standard to evaluate the cellular respiration within OptoElecWell device. After 15 minutes of basal respiration monitoring, we injected known modifiers of cellular oxygen consumption: FCCP (1µM), an uncoupler of oxidative phosphorylation that increases the cellular oxygen consumption, followed by the addition of metabolic poison Antimycin A (10 µM), a specific inhibitor of mitochondrial electron transport chain. Figure 2c, shows the variations of cellular oxygen uptake with respect to the additions of FCCP, where there is an increase of the rate of oxygen consumption (1.5 µA/min) compared to the basal respiration of cells (0.6 µA/min) and the effect of drug Antimycin A, i.e., impairment of oxygen consumption. In order to demonstrate the effect of these drugs specifically towards the cells within the microwell array, the same experiment was repeated by adding the metabolic modulators in inverse fashion, i.e., addition of Antimycin A first, followed by FCCP and we observed no significant increase of oxygen uptake after the addition of FCCP.

Overall, the results obtained by monitoring oxygen consumption of cells, were fully consistent with the results widely reported using different technologies and well understood in conditions of modulations of the respiratory chain activity (Hill et al., 2012; Ruas et al., 2016; Santos et al., 2017; Simonnet et al., 2014). However, the higher interest of this presented OptoElecWell approach resides in the simultaneous use of high-resolution fluorescence imaging and electrochemical sensing which eventually leads to the dynamic monitoring of at least two-cell metabolic parameters (biophysical and biochemical properties). In this context, we monitored the effect of the concentrations of the carbon source (glucose) on cellular oxygen uptake rate linked with OXPHOS metabolism and the intracellular rheological properties within the cell, known to be impacted by metabolic changes.

### Dual detection: the effect of glucose concentration on the basal oxygen uptake rate and macromolecular diffusion

Glucose is an essential carbon source for the production of ATP. The latter is mainly synthesized through glycolysis and oxidative phosphorylation through the mitochondrial respiration. It was recently shown that modulation of intracellular ATP concentration affected microrheological properties: lower ATP concentration decreased the diffusion of tracer particles (Mathieu and Manneville, 2019). It is critical for cells to sense and adapt to potential changes in glucose concentration. Yeast (*S. cerevisiae)* is as an excellent model organism to study glucose-derived metabolism through mitochondrial respiration (Conrad et al., 2014; Galdieri et al., 2010). Yeasts were cultured in media containing different glucose concentrations ranging from 0.5% to 2% of glucose. The media was osmotically compensated for the lower glucose concentrations (0.5% and 1%) in order to avoid additional osmotic perturbations in comparison with the control (2%). The effect of extracellular concentration of the glucose on both mitochondrial specific oxygen consumption rate (basal respiration) and internal dynamics of molecular diffusion was simultaneously measured on OptoElecWell device (figures 3 and 4).

**Figure 3:**
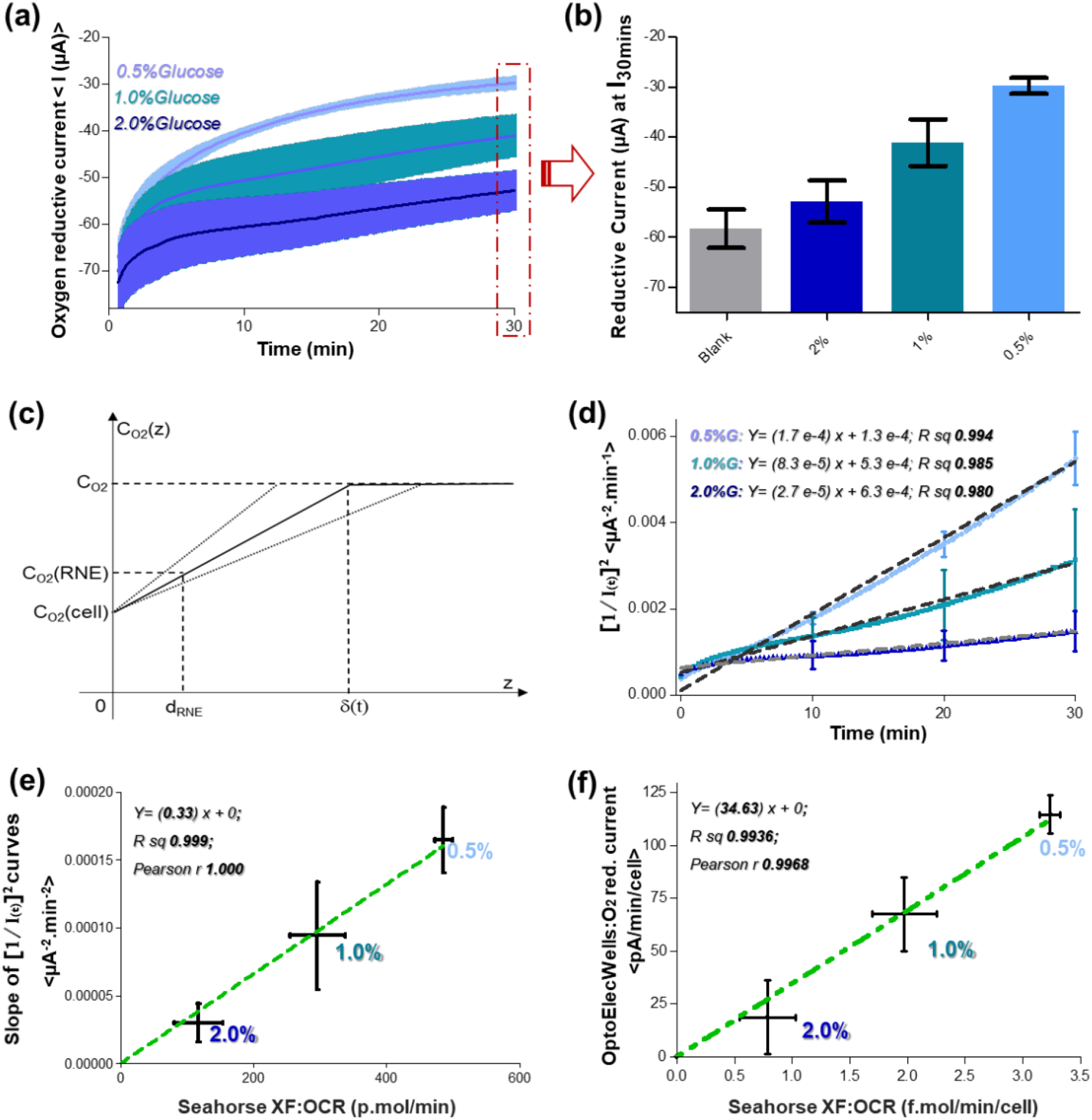
The effect of extracellular concentration of glucose on mitochondria specific cellular basal oxygen uptake rate: (a) Kinetic evolution and (b) end point values, at 30 mins, of the oxygen reductive peak current biased at −0.7 V Vs Ag/AgCl measured on each CV scan, under different substrate concentrations ranging from 0.5% to 2% of glucose. The standard deviations of each condition were represented as shaded error bands in the case of kinetic evolution; (c) Theoretical model for OptoElecWell based cellular oxygen consumption monitoring and the effect of substrate concentration on their oxygen consumption kinetics. C_O2_, C_O2_(RNE) and C_O2_(cell) are the dissolved oxygen concentrations in the bulk solution, at the ring nanoelectrode surface and at the cell surface, d_RNE_ is the distance between ring nanoelectrode and the cell, and δ(t) is the time-dependent diffusion layer thickness; (d) Time-evolution of inverse square peak current values at different glucose concentrations showing quasi-linear variations, where I(t) represents i_final_ – i_measured_; (d) The slopes of inverse square peak current values at different glucose concentrations were compared to the responses obtained from a commercial Seahorse XF. The number of cells used for cellular respiration measurement in OptoElecWell device and SeaHorse XF were approximately 8.0 x 10^3^ and 1.5 x 10^5^, respectively. The Pearson correlation and R-square values were determined to be 1.000 and 0.999, respectively where n=3. (e) Oxygen reductive current values (blank subtracted) of OptoElecWells and Seahorse XF OCR readings were normalized, per cell, per minute, to relate the obtained reductive current values to cellular oxygen consumption rates. The Pearson correlation and R-square values were determined to be 0.9968 and 0.9936, respectively where n=3.

**Figure 4:**
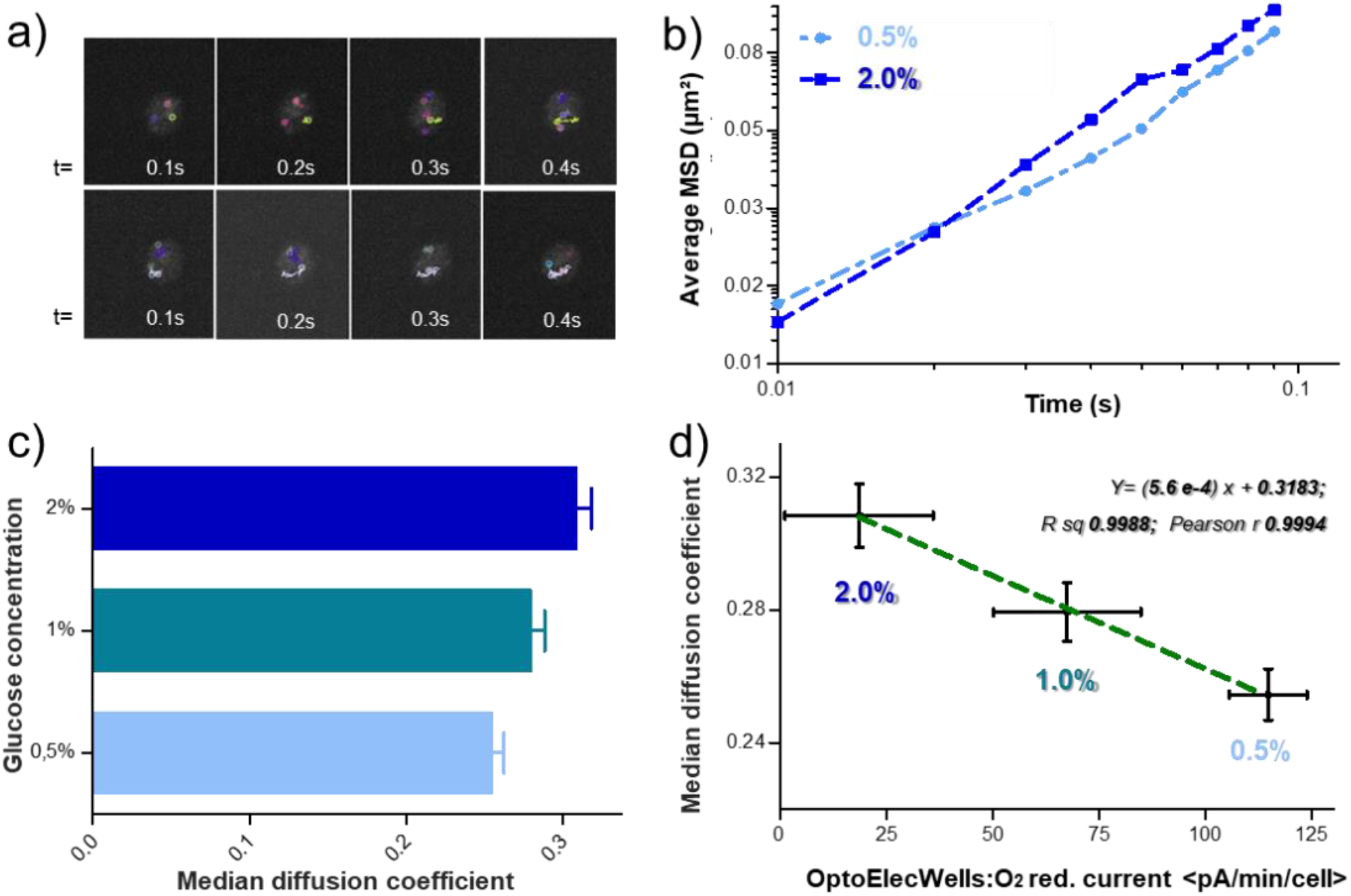
(a) Time-lapse single particle tracking of entrapped yeast cells within the microwells. 40nm GEMs were visualized using the T-sapphire fluorescent protein (green) and the diffusive trajectories of each particle were monitored using inverted confocal microscope (63x objective). (b) The resultant average MSD versus time plot of entrapped yeast cells grown at 0.5% and 2% glucose concentration; (c) The effect of extracellular concentration of glucose on the effective diffusion coefficients in yeast cells; (d) Evaluation of the relation between molecular diffusion and cellular basal oxygen respiration using OptoElecWells: The normalized oxygen reductive peak current values, per minute, per cell, at different glucose concentrations were compared to the responses obtained from GEMs mediated molecular diffusion coefficient measurements. The R-square and Pearson correlation values were determined to be 0.9988 and 0.9994, respectively (n=3).

### Basal cellular respiration monitoring at different glucose concentrations

Thanks to the high density of ring-shaped nanoelectrodes located within the microwells and the continuous CV-based sensing methodology, basal cellular oxygen uptake kinetics (figure 3a) and end-point measurements (figure 3b) were performed. The measured oxygen reduction current values, with respect to time, suggested that the oxygen uptake rate is at its maximum when the cells were exposed to 0.5% glucose (steep curve). Further increase in the extracellular glucose concentration resulted in a progressive decrement in the oxygen reduction peak currents, suggesting inferior oxygen uptake. However, the basal respiration monitoring at different glucose concentrations are trickier to interpret than the respiration measurements under metabolic modulators (figure 2c-d). In the latter case, the respective oxygen consumption slopes of different activators and inhibitors were directly compared to the basal respiration within the same experiment. Hence, we proposed a theoretical model to comprehensively understand the OptoElecWell-based cellular oxygen consumption monitoring and the effect of substrate (glucose) concentration on their oxygen consumption kinetics (see Methods section and figure 3c). As shown in the figure 3d (see also Methods), we estimated I(t) [(i_measured_ - i_final_)] values for different glucose concentrations and the evolution of inverse square currents [1/I(t)]^2^ was plotted with respect to time. In all three cases, quasi-linear variations were effectively obtained, demonstrating the “Cottrell-like behaviour” in terms of cellular oxygen consumption. To further validate our methodology, the measured slopes of inverse square peak current values [1/I(t)]^2^ at different glucose concentrations using OptoElecWell device were compared with the OCR values obtained by a commercial gold standard SeaHorse XF respirometer. We observed an excellent positive correlation between the two technologies, as demonstrated with the Pearson correlation and R-square values of 1.000 and 0.999 respectively (figure 3e).

In addition, we also normalized the readings of both techniques, per cell, per minute, in an attempt to relate the obtained reductive current “I_30mins_” values (blank subtracted) to the cellular oxygen consumption rates (figure 3f). Again we observed an excellent positive correlation and found out that a reductive current change of 100 pA was equivalent to approximately 3 fM of basal cellular oxygen consumption. Even though the variability of measurements in the OptoElecWell device is on par with the Seahorse XF analyser (Pearson correlation and R-square values of 0.9968 and 0.9936 respectively), our device can function with at least 20 times lesser biological sample (10^3^ to 10^4^ cells) than the Seahorse XF respirometer (1.5 × 10^5^ cells) and in addition, provides an optical means to monitor the contents within single cells in a high-throughput manner.

### Optical monitoring of molecular diffusion through single-particle tracking

The great optical capabilities of our OptoElecWell device allow for high-throughput confocal imaging inside cells (figure 2d). We imaged GEMs nanoparticles in cells using a spinning disk confocal microscope at 50 Hz imaging rate (see Methods). Single-particle tracking is possible in these conditions (figure 4a). Note that the imaging was performed while oxygen was electrochemically monitored in the device. We analysed the trajectories as in (Delarue et al., 2018) in order to extract ensemble-averaged mean-square displacement (MSD) curves and single particle short-term diffusion coefficients (figure 4b-c). We observed that particles in lower glucose concentration diffused slower, with up to 20% increase between 0.5% and 2% glucose. Interestingly, diffusion seems to become anomalous under low oxygen concentration, with the power exponent of the MSD lower than 1. We plotted the measured diffusion coefficient, reflecting the viscosity of the cell interior, as a function of the oxygen consumption rate measured electrochemically. We observed an anti-correlation of the two quantities (figure 4d): diffusion increases as oxygen consumption rate decreases (Pearson correlation and R-square values of 0.9988 and 0.9994 respectively). While an observed decrease in oxygen respiration rates (aerobic metabolism) at higher glucose concentrations has been observed in the past, its link with changes in rheological parameters has never been established before.

### Conclusion

We demonstrated in this article the feasibility of dual detection of electrochemical readout (oxygen here) and high-resolution fluorescence imaging (SPT of fluorescent nanoparticles). Our Oxygen detection technique is at the level of “state-of-the-art” in terms of accuracy and requires lower amounts of cellular material, compared to the commercial gold standard. As a proof of concept, we explored the understudied link between cellular metabolism and intracellular rheological properties. We find that while respiration increases, the diffusion of tracer particles decreases with lower oxygen diffusion. Respiration is linked with ATP production through mitochondria, and it is known that decreased biochemical activity associated with ATP depletion dramatically decreases tracers diffusion (Munder M et al., 2016). Our result thus appears as counter-intuitive: while respiration increases, ATP production is expected to increase, with a potential increase in biochemical activity, which should fluidize the cytoplasm. We find the opposite: an increased respiration correlates with a decreased diffusion.

This result calls for a better understanding of the link between metabolism and intracellular rheology. Devices such as OptoElecWell are ideal to conduct these studies, coupling electrochemistry and high-resolution optics. In the present study and as a proof of concept, we only addressed the link between oxygen consumption rates with cellular rheology. We believe that this methodology serves as a starting point to further explore the complex relations between the cellular metabolic and rheological indicators. Because the platform described here has been designed to be flexible to allow higher levels of multiplexing, analytes such as glucose or lactate can be sensed along with other metabolic and rheological parameters with some minor surface modifications to the electrode surfaces (Soldà A et al., 2017; Zhang et al., 2012). Also note that the electrodes can function to clamp some physical parameters, such as oxygen: one electrode can be used to consume oxygen and impose a given concentration to infer the impact of different oxygen concentrations on metabolism and intracellular rheological properties.

Overall, our novel generation of OptoElecWell with dual detection approach takes simple cell-based analyses to multidimensional, multiparametric levels. It represents a powerful analytical tool, allowing rapid, simultaneous opto-electrochemical analyses of a large number of cells, while retaining the ability to address the single cells within the array. This study provides a base that can enable the exploration of the missing links between the cellular metabolism (or cellular physiological responses) and its intracellular (rheological) properties.

